# Experimental analysis of exome-scale mutational signature of glycidamide, the reactive metabolite of acrylamide

**DOI:** 10.1101/254664

**Authors:** Maria Zhivagui, Maude Ardin, Alvin W.T. Ng, Mona I. Churchwell, Manuraj Pandey, Stephanie Villar, Vincent Cahais, Alexis Robitaille, Liacine Bouaoun, Adriana Heguy, Kathryn Guyton, Martha R. Stampfer, James McKay, Monica Hollstein, Magali Olivier, Steven G. Rozen, Frederick A. Beland, Michael Korenjak, Jiri Zavadil

**Affiliations:** Molecular Mechanisms and Biomarkers Group, International Agency for Research on Cancer, Lyon 69008, France; Centre for Computational Biology, Duke-NUS Medical School, Singapore 169857, Singapore; Program in Cancer and Stem Cell Biology, Duke-NUS Medical School, 169857, Singapore; NUS Graduate School for Integrative Sciences and Engineering, 117456, Singapore; Division of Biochemical Toxicology, National Center for Toxicological Research, Jefferson, AR 72079, USA; Epigenetics Group, International Agency for Research on Cancer, Lyon 69008, France; Infections and Cancer Biology Group, International Agency for Research on Cancer, Lyon 69008, France; Environment and Radiation Section, International Agency for Research on Cancer, Lyon 69008, France; Department of Pathology and Genome Technology Center, New York University, Langone Medical Center, New York, NY 10016, USA; IARC Monographs Section, International Agency for Research on Cancer, Lyon 69008, France; Biological Systems and Engineering Division, Lawrence Berkeley National Laboratory, Berkeley, CA, 94720, USA; Genetic Cancer Susceptibility Group, International Agency for Research on Cancer, Lyon 69008, France; Deutsches Krebsforschungszentrum, 69120 Heidelberg, Germany; Faculty of Medicine and Health, University of Leeds, LIGHT Laboratories, Leeds LS2 9JT, United Kingdom

**Keywords:** Acrylamide, glycidamide, DNA adducts, massively parallel sequencing, mutational signatures

## Abstract

Acrylamide, a probable human carcinogen, is ubiquitously present in the human environment, with sources including heated starchy foods, coffee and cigarette smoke. Humans are also exposed to acrylamide occupationally. Acrylamide is genotoxic, inducing gene mutations and chromosomal aberrations in various experimental settings. Covalent haemoglobin adducts were reported in acrylamide-exposed humans and DNA adducts in experimental systems. The carcinogenicity of acrylamide has been attributed to the effects of glycidamide, its reactive and mutagenic metabolite capable of inducing rodent tumors at various anatomical sites. In order to characterize the pre-mutagenic DNA lesions and global mutation spectra induced by acrylamide and glycidamide, we combined DNA-adduct and whole-exome sequencing analyses in an established exposure-clonal immortalization system based on mouse embryonic fibroblasts. Sequencing and computational analysis revealed a unique mutational signature of glycidamide, characterized by predominant T:A>A:T transversions, followed by T:A>C:G and C:G>A:T mutations exhibiting specific trinucleotide contexts and significant transcription strand bias. Computational interrogation of human cancer genome sequencing data indicated that a combination of the glycidamide signature and an experimental benzo[*a*]pyrene signature are nearly equivalent to the COSMIC tobacco-smoking related signature 4 in lung adenocarcinomas and squamous cell carcinomas. We found a more variable relationship between the glycidamide‐ and benzo[*a*]pyrene-signatures and COSMIC signature 4 in liver cancer, indicating more complex exposures in the liver. Our study demonstrates that the controlled experimental characterization of specific genetic damage associated with glycidamide exposure facilitates identifying corresponding patterns in cancer genome data, thereby underscoring how mutation signature laboratory experimentation contributes to the elucidation of cancer causation.

**A 40-word summary:** Innovative experimental approaches identify a novel mutational signature of glycidamide, a metabolite of the probable human carcinogen acrylamide. The results may elucidate the cancer risks associated with exposure to acrylamide, commonly found in tobacco smoke, thermally processed foods and beverages.

## Introduction

Cancer can be caused by chemicals, complex mixtures, occupational exposures, physical agents, and biological agents, as well as lifestyle factors. Many human carcinogens show a number of characteristics that are shared among carcinogenic agents (1). Different human carcinogens may exhibit a spectrum of these key characteristics, and operate through separate mechanisms to generate patterns of genetic alterations. Recognizable patterns of genetic alterations or mutational signatures characterize carcinogens that are genotoxic. Recent work shows that these DNA sequence changes can be expressed in simple mathematical terms that enable mutational signatures to be extracted from thousands of cancer genome sequencing data sets (2). Several of the over 30 identified mutational signatures have been attributed to specific external exposures or endogenous factors through epidemiological and experimental studies (2). However, about 40% of the current signatures remain of unknown origin, and additional, thus far unrecognized, signatures are likely to be defined in rapidly accumulating cancer genome data. Well-controlled experimental exposure systems can thus help identify the underlying causes of known orphan mutational signatures as well as define new patterns generated by candidate carcinogens (reviewed in (3,4)).

Various diet-related exposures contribute to the human cancer burden. Examples include contaminants in food or alternative medicines, such as aflatoxin B1 (AFB1) or aristolochic acid (AA). The mutagenicity of these compounds is well-documented; AFB1 induces predominantly C:G>A:T base substitutions and AA causes T:A>A:T transversions. The characteristic mutations coupled with information on the preferred sequence contexts in which they are likely to arise allowed unequivocal association of exposure to AFB1 or AA with specific subtypes of hepatobiliary or urological cancers, respectively (5-13).

Among dietary compounds with carcinogenic potential, acrylamide is of special interest due to extensive human exposure. Important sources of exposure to acrylamide include tobacco smoke (14), coffee (15), and a broad spectrum of occupational settings (16). Dietary sources of acrylamide comprise carbohydrate-rich food products that have been subject to heating at high temperatures. This is due to Maillard reactions, which involve reducing sugars and the amino acid asparagine, present in potatoes and cereals (17). There is sufficient evidence that acrylamide is carcinogenic in experimental animals (18,19) and it has been classified as a probable carcinogen (Group 2A) by the International Agency for Research on Cancer in 1994 (16). The association of dietary acrylamide exposure with renal, endometrial and ovarian cancers has been explored in recent epidemiological studies (20,21). However, accurate acrylamide exposure assessment in epidemiological studies based on questionnaires has been difficult, and more direct measures of molecular markers, such as hemoglobin adduct levels, may not yield conclusive findings on past exposures (22-27). An improved understanding of its mechanism of action using well-controlled experimental systems is critical for understanding the potential carcinogenic risk associated with exposure.

Acrylamide undergoes oxidation by cytochrome P450, producing the reactive metabolite glycidamide that is highly efficient in DNA binding due to its electrophilic epoxide structure (28-30). The *Hras* mutation load in neoplasms of mice exposed to acrylamide or glycidamide was found to be considerably higher in mice treated with glycidamide (31). This finding is corroborated by a considerably higher mutation frequency in the *cII* reporter gene of Big Blue mouse embryonic fibroblasts treated with glycidamide in comparison to acrylamide (32,33). Mutation analysis in different experimental *in vivo* and *in vitro* models using reporter genes showed an increased association of acrylamide and glycidamide exposure with T:A>C:G transitions, as well as T:A>A:T and C:G>G:C transversion mutations (31-36), whereas glycidamide exposure was also characterized by C:G>A:T transversions (33). However, these proposed acrylamide‐ and glycidamide-specific mutation patterns were based on limited mutation counts in reporter genes and thus do not reflect the complexity of genome-wide distributions and profiles. Based on the limited data available thus far, it is not possible to translate adequately the reported mutation types (T:A>C:G, T:A>A:T, C:G>G:C, C:G>A:T) to global alteration patterns.

The advent of massively parallel sequencing has created the opportunity to study a large number of mutations in a single sample, thus significantly enhancing the power of mutation analysis in experimental models and enabling reliable identification of specific sequence contexts for the induced alterations. Analogously to human cancer genome projects, genome-scale mutational signatures can be extracted from highly controlled carcinogen exposure experiments using mammalian cell and animal models coupled with advanced mathematical approaches (2,3,37,38).

Here we report the systematic assessment of acrylamide and glycidamide mutagenicity based on DNA adduct formation and mutation profile analysis using massively parallel sequencing in a cell model amenable to the analysis of carcinogen-induced mutation patterns and their impact on the resulting cell phenotype (3,37-39). We identify a specific and robust mutational signature attributable to glycidamide, and by computationally interrogating human cancer genome-wide mutation data, we characterize glycidamide signature-positive tumors, thereby highlighting a potential contribution of acrylamide/glycidamide exposure to carcinogenesis in humans.

## Materials and methods

### Source and authentication of primary cells

Primary <u>Hu</u>man-<u>p</u>53 <u>k</u>nock-<u>i</u>n mouse embryonic fibroblasts (Hupki MEFs) were isolated from 13.5-day old *Trp53^tm/Holl^* mouse embryos from the Central Animal Laboratory of the Deutsches Krebsforschungszentrum, Heidelberg, as described previously (40). The mice had been tested for Specific Pathogen-Free (SPF) status. The derived primary cells were genotyped for the human *TP53* codon 72 polymorphism (Table 1) to authenticate the embryo of origin. Cells from three different embryos (E210, E213 and E214) were used for the exposure experiments (Table 1). All subsequent cell cultures were routinely tested at all stages for the absence of mycoplasma.

**Table 1:** Summary of cell lines, treatment conditions and *TP53*^1^ mutation status.

| Sample ID | Embryo | Exposure | Conc. (mM) | Exposure duration (hrs) | coding DNA change<sup>2</sup> | genomic DNA change<sup>3</sup> | aa change | Codon 72 (rs1042522)<sup>4</sup> |
| --- | --- | --- | --- | --- | --- | --- | --- | --- |
| Prim_1 | E210 | - | - | - |  |  |  | Pro/Pro |
| Prim_2 | E213 | - | - | - |  |  |  | Arg/Pro |
| Prim_3 | E214 | - | - | - |  |  |  | Pro/Pro |
| Spont_1 | E213 | - | - | - |  |  |  | Arg/Pro |
| Spont_2 | E214 | - | - | - |  |  |  | Pro/Pro |
| Spont_3 | E214 | - | - | - |  |  |  | Pro/Pro |
| ACR_S9_1 | E213 | ACR | 5 | 24 |  |  |  | Arg/Pro |
| ACR_S9_2 | E213 | ACR | 5 | 24 |  |  |  | Arg/Pro |
| ACR_1 | E213 | ACR | 10 | 24 | c.881delA | g.7577057delT | p.E294fs | Arg/- |
| ACR_2 | E213 | ACR | 10 | 24 | c.818G>T | g.7577120C>A | p.R273L | Pro/- |
| ACR_3 | E214 | ACR | 10 | 24 | c.740A>T; c.839G>C | g.7577541T>A; g.7577099C>G | p.N247I; p.R280T | Pro/Pro |
| ACR_4 | E214 | ACR | 10 | 24 |  |  |  | Pro/Pro |
| ACR_5 | E214 | ACR | 10 | 24 |  |  |  | Pro/Pro |
| GA_1 | E210 | GA | 3 | 24 |  |  |  | Pro/Pro |
| GA_2 | E210 | GA | 3 | 24 |  |  |  | Pro/Pro |
| GA_3 | E210 | GA | 3 | 24 | c.309-310CC>TA | g.7579377-7579378GG>TA | [p.Y103Y; p.Q104K] | Pro/Pro |
| GA_4 | E214 | GA | 3 | 24 |  |  |  | Pro/Pro |
| GA_5 | E214 | GA | 3 | 24 |  |  |  | Pro/Pro |

<sup>1</sup> human TP53 gene; <sup>2</sup> NM\_000546.4 coding sequence; <sup>3</sup> hg19 genomic coordinates; <sup>4</sup> human polymorphic site (rs1042522)
Prim = Primary cells; Spont = spontaneously immortalized clones; ACR = acrylamide-exposure derived clones; GA = glycidamide-exposure derived clones. Each exposure condition was carried out in two biological replicates (embryos). S9 = human S9 fraction; Pro = proline; Arg = arginine; Arg/- or Pro/- = loss of allele; fs = frameshift; aa = amino acid.

### Cell culture, exposure and immortalization

The primary MEF cells were expanded in Advanced DMEM supplemented with 15% fetal calf serum, 1% penicillin/streptomycin, 1% pyruvate, 1% glutamine, and 0.1% p-mercapto-ethanol. The cells were then seeded in six-well plates and, at passage 2, exposed for 24 hours to acrylamide (A4058, Sigma), glycidamide (04704, Sigma), or vehicle (PBS). Acrylamide exposure was carried out in the absence or presence of 2% human S9 fraction (Life Technologies) complemented with NADPH (Sigma). Exposed and control primary cells were cultivated until they bypassed senescence and immortalized clonal cell populations could be isolated (41). The human mammary epithelial cell (HMEC) cultures utilized in this study for whole-genome sequencing (WGS) were generated from benzo[*a*]pyrene (B[a]P) exposed HMEC described previously (42,43).

### MTT assay for cell metabolic activity and viability

Cells were seeded in 96-well plates and treated as indicated. Cell viability was measured 48 hours after treatment cessation using CellTiter 96^®^ Aqueous One solution Cell Proliferation Assay (Promega). Plates were incubated for 4 hours at 37°C and absorbance was measured at 492 nm using the APOLLO 11 LB913 plate reader. The MTT assay was performed in triplicates for each experimental condition.

### γH2Ax Immunofluorescence

Immunofluorescence staining was carried out using an antibody specific for Ser139-phosphorylated H2Ax (γH2Ax) (9718, Cell Signaling Technology). Primary MEFs were seeded on coverslips in 12 well-plates. The cells were incubated in with γH2Ax-antibody (1:500 in 1% BSA) at 4°C overnight. Subsequent incubation with a fluorochrome-conjugated secondary antibody (4412, Cell Signaling Technology) was carried out for 60 minutes at room temperature. Coverslips were mounted in Vectashield mounting medium with DAPI (Eurobio). Immunofluorescence images were captured using a Nikon Eclipse Ti.

### DNA adduct analysis

Glycidamide-DNA adducts (N7-(2-carbamoy-2-hydroxyethyl)-guanine (N7-GA-Gua) and N3-(2-carbamoy-2-hydroxyethyl)-adenine (N3-GA-Ade)) were quantified by liquid chromatography-mass spectrometry (LC-MS/MS) with stable isotope dilution as previously described (44) (see Supplementary Materials and Methods for details). The LC-MS/MS used for quantification consisted of an Acquity UPLC system (Waters) and a Xevo TQ-S triple quadrupole mass spectrometer (Waters). The same MRM transitions as previously described (44) were monitored with a cone voltage of 50V and collision energy of 20eV for each adduct transition and its corresponding labeled isotope transition.

### TP53 genotyping

Exons 4 to 8 of the knocked-in human *TP53* gene (NC_000017.11) were sequenced using standard protocols. Sanger sequencing of PCR products was performed at Biofidal (Lyon, France). *TP53* primer sequences are listed in Supplementary Materials and Methods. Resulting sequences were analyzed using the CodonCode Aligner software.

### Library preparation and whole-exome sequencing (WES)

Library preparation was carried out using the Kapa Hyper Plus library preparation kit (Kapa Biosystems) according the manufacturer’s instructions. Exome capture was performed using the SureSelect XT Mouse All Exon Kit (Agilent Technologies). Eighteen exome-captured libraries were sequenced in the paired-end 150 base-pair run mode using the Illumina HiSeq4000 sequencer.

### Processing of WES data

Fastq files were analyzed for data amount and quality using FastQC (0.11.3) and were processed with an in-house pipeline for adapter trimming and alignment to the mm10 genome (release GRCm38). These components of the pipeline are publicly available at https://github.com/IARCbioinfo/alignment-nf. The resulting alignment files had a mean depth-of-coverage of 135 and 175 for acrylamide and glycidamide samples, respectively. All alignment files can be accessed from the NCBI Sequence Read Archive (SRA) data portal under the BioProject accession number PRJNA238303. Two somatic variant callers were employed with default parameters in order to detect single base substitutions (SBS) and small insertions/deletions (indels) (MuTect 1.1.6-4 and Strelka 1.015) in exposed clones, using primary cells as normal samples. Each immortalized clone was compared to primary MEFs from three different embryos (conditions Prim_1, Prim_2, and Prim_3). The overlap of the variant calling outcome with respect to the different primary MEFs showed concordance close to 80% (Suppl. Fig. S1) with MuTect exhibiting more stringent calling performance. Thus, mutation data obtained from the MuTect variant caller were further processed with the MutSpec suite ((45); https://github.com/IARCbioinfo/mutspec). For more details, see Supplementary Materials and Methods and the summary of sequencing metrics (Suppl. Table S1 ‐ not available in the preprint version), the list of identified MuTect SBS variants (Suppl. Table S2 – not available in the preprint version) and indels (Suppl. Table S3 ‐ not available in the preprint version).

### Bioinformatics and statistical analyses

The FactoMiner R package (R package version 3.3.2; https://cran.r-project.org/web/packages/FactoMineR) was used to perform the principal component analysis (PCA). To perform the transcription strand bias (SB) analyses, *p*-values were calculated using Pearson’s *χ*^2^ test. As multiple comparisons were assessed, the *p*-value was adjusted by applying a false discovery rate (FDR). Statistical analyses were carried out using the stats R package. The SB was considered statistically significant at *p*-value ≤ 0.05. To analyze samples mutation spectra and treatment-specific mutational signatures, filtered mutations were classified into 96 types corresponding to the six possible base substitutions (C:G>A:T, C:G>G:C, C:G>T:A, T:A>A:T, T:A>C:G, T:A>G:C) and the 16 combinations of flanking nucleotides immediately 5’ and 3’ of the mutated base. Mutation patterns were then deconvoluted into mutational signatures using the non-negative matrix factorization (NMF) algorithm (46,47). The reconstruction error calculation evaluated the accuracy with which the deciphered mutational signatures describe the original mutation spectra of each sample by applying Pearson correlation and cosine similarity.

In order to clean up the profile of the glycidamide mutational signature from the residual signature 17 signal and to increase the stability of NMF decomposition, we supplied the NMF input by adding samples with a high level of signature 17 (over 65% contribution as determined by independent NMF analysis, see Supplementary Materials and Methods).

Cosine similarity analysis was used to evaluate the concordance of the newly identified T:A>A:T-rich mutational signature of glycidamide with the previously reported mutational signatures characterized by a predominant T:A>A:T content. These comprised COSMIC signatures 22 (AA), 25 and 27 (both of unknown etiology(2)), the experimentally derived mutational signature of AA (37,45), 7,12-dimethylbenz[*a*]anthracene (DMBA) (48,49), and urethane (50).

We employed the mutational signature activity (mSigAct) software’s sparse signature assignment function (sparse.assign.activity) (13) to assess the presence of the experimental mutational signatures of glycidamide and benzo[*a*]pyrene in whole-genome somatic mutation data from 38 lung adenocarcinomas, 48 lung squamous carcinomas, and 320 liver cancers from the ICGC Pan-Cancer Analysis of Whole Genomes (PCAWG) study. We excluded 244 hyper-mutated microsatellite unstable and aristolochic acid signature-containing liver tumors as the presence of high numbers of T>A mutations adversely prevented assessment of the possible presence of the glycidamide signature. A set of 11 active COSMIC mutational signatures were identified in the remaining tumor samples (excluding COSMIC signature 4).

We defined a ‘pure’ experimental C>N benzo[*a*]pyrene signature by WGS (using Illumina HiSeq4000 by Genewiz, NJ, USA) of finite lifespan post-stasis clones derived from primary human mammary epithelial cells (HMEC) treated with B[a]P as previously described (42,43,51). The read alignment to NCBI GRCh38 genome build, variant calling, filtering and annotation were consistent with the MutSpec pipeline described above (45). Proportion matrices of the experimental GA-signature, the GA-signature normalized to the human genome trinucleotide frequency to allow for human PCAWG data screening, and the whole-genome B[a]P signature are available in Suppl. Table S4 (not accessible in the preprint version).

## Results

### Acrylamide and glycidamide induce cytotoxic and genotoxic responses in Hupki MEFs

Upon exposure of primary Hupki MEFs to a range of concentrations of acrylamide (ACR) (in the absence or presence of the S9 fraction) and its metabolite, glycidamide (GA), we observed a dose-dependent cytotoxic effect on the cells for either compound (Fig. 1A). This analysis informed the selection of two conditions for the ACR exposure to be used in the subsequent exposure/immortalization experiments, 10 mM ACR for 24 hours in the absence of human S9 fraction, and 5 mM ACR for 24 hours in the presence of S9 fraction, which elicited 50% (range 30-70%) decrease in cell viability. The IC50 condition for GA was used for subsequent mutagenesis analysis, corresponding to a 24-hour treatment with 3 mM of the compound. The genotoxic effects of either ACR or GA manifested by a marked increase in γH2Ax staining in the exposed cell populations, in comparison to the mock-treated control cells (Fig. 1B).

**Figure 1:**
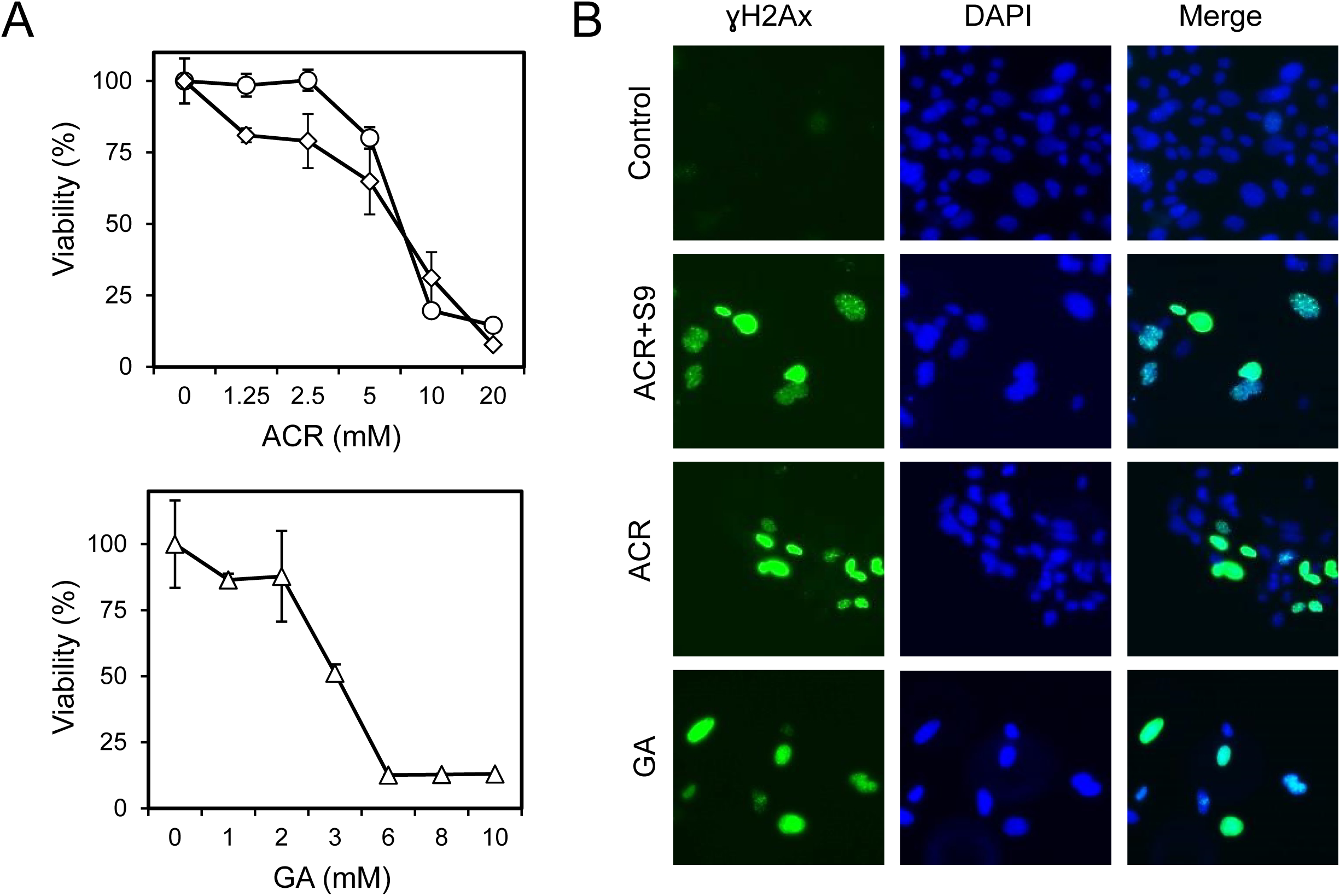
Acrylamide‐ and glycidamide-induced cytotoxicity and genotoxicity *in vitro*. (**A**) Cell viability, following 24-hour treatment of primary MEFs with the indicated concentrations of acrylamide (top panel), in the absence (diamonds) and presence (circles) of human S9 fraction, and glycidamide (bottom panel), as determined by MTT assay. Absorbance was measured 48 hours after treatment cessation and was normalized to untreated cells. The results are expressed as mean percent ±SD of three replicates. (**B**) DNA damage assessment by immunofluorescence with an antibody specific for Ser139-phosphorylated histone H2Ax (?H2Ax). Primary MEFs were treated with acrylamide or glycidamide for 24 hours prior to immunofluorescence. Compound concentrations used were based on 20-70% viability reduction in the MTT assay: 10 mM acrylamide, 5 mM acrylamide in the presence of S9 fraction and 3 mM glycidamide. ACR: acrylamide; GA: glycidamide.

### Immortalized MEF cells accumulate *TP53* mutations following acrylamide or glycidamide treatment

Primary MEF cultures from three different embryos (Prim_1, Prim_2, and Prim_3) were exposed to ACR or GA using the established conditions and multiple immortalized clones were derived. MEF senescence and immortalization phases were evident from the growth curves generated for each culture (Suppl. Fig. S2). Subsequently, the clones derived from ACR exposure (ACR clones) and GA exposure (GA clones) and spontaneous immortalization (Spont), were pre-screened for *TP53* mutations by Sanger sequencing, to assess the mutagenic process prior to exome-scale analysis. In the context of ACR treatment, clones obtained from the Prim_2 MEFs that were heterozygous for the polymorphic site in codon 72 showed a loss of heterozygosity involving a loss of the proline allele in the ACR_1 clone whereas the arginine allele was lost in ACR_2, giving rise to a hemizygous clone (Table 1). No *TP53* mutations were observed in any of the three Spont clones, whereas 3 out of 7 ACR clones and 1 of 5 GA clones carried non-synonymous *TP53* mutations (Table 1). The detected mutations indicated specific selection for mutations in the *TP53* gene during cell immortalization and confirmed the clonal nature of MEF immortalization.

### Analysis of mutation spectra

Whole-exome sequencing of all spontaneously immortalized and exposed clones and subsequent extraction of acquired variants revealed that the total number of acquired SBS did not differ markedly between the ACR and Spont clones. The Spont clones harbored on average 190 (median = 151, range = 141-277) SBS, whereas the ACR clones had on average 208 (median = 173, range = 151-262) SBS. In contrast, the total number of SBS was considerably increased in the GA clones, with an average of 485 SBS (median = 448, range = 370-592) (Suppl. Table S1 and S2 – not available in the preprint version). This finding suggests markedly stronger mutagenic properties of GA in the MEFs. To estimate the extent of sequencing-related damage in our samples, we determined the GIV score of each sample as described in Materials and Methods and in (52). No detectable damage for any of the mutation types was observed in our dataset (data not shown). The ACR exposed samples exhibited an overall diffuse pattern across the six different SBS types (Suppl. Fig. S3). The Spont clones showed an enrichment of C:G>G:C SBS in the 5’-GCC-3’ context, which was also present at varying levels in the exposed cultures. This particular mutation type appears to be related to the culture conditions used for the immortalization assay, as its presence has previously been noted upon spontaneous as well as exposure-driven MEF immortalization (37). No significant transcription strand bias was observed for any of the mutation classes in the Spont or ACR clones (Suppl. Fig. S4). In the five clones derived from the GA-treated primary MEF cultures, we observed an enrichment of acquired T:A>A:T and C:G>A:T transversions and T:A>C:G transitions (Suppl. Fig. S3B), marked by significant transcription strand bias (Suppl. Fig. S4).

PCA performed on the resulting 6-class SBS spectra unambiguously separated the GA clones from the remaining experimental conditions (Fig. 2A). The analysis of indels (listed in Suppl. Table S3 – not available in the preprint version) showed lower numbers of these alterations in the GA-associated clones compared to the ACR or Spont clones (Fig. 2B). This suggests that a higher accumulation of SBS may selectively promote the senescence bypass and selection of the GA clones, with a decreased functional contribution of indels, while an inverse scenario is plausible in case of the Spont and ACR clones, reminiscent of a previous report based on the Big Blue mouse embryonic fibroblasts and c*II* transgene (53).

**Figure 2:**
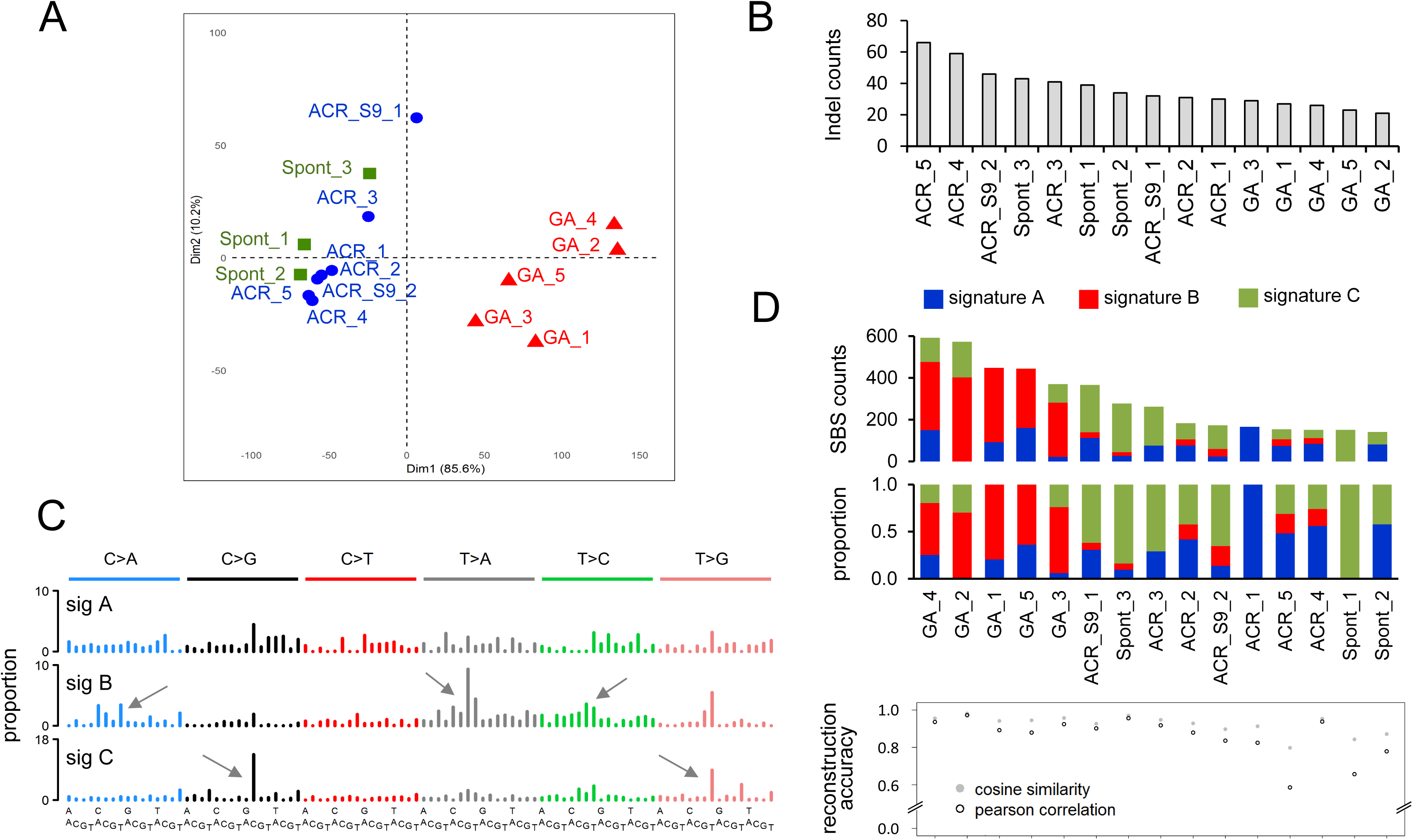
Analysis of the mutation patterns derived from exome sequencing data from immortalized Hupki MEF clones. (**A**) Principle component analysis (PCA) of WES data. PCA was computed using as input the mutation count matrix of the clones that immortalized spontaneously (Spont) or were derived from exposure to acrylamide (ACR) or glycidamide (GA). Each sample is plotted considering the value of the first and second principal components (Dim1 and Dim2). The percentage of variance explained by each component is indicated within brackets on each axis. Spont, ACR‐ and GA-exposed samples are represented by differently colored symbols. (**B**) Representation of small insertions and deletions (indels) counts within the immortalized clones as determined by the Strelka variant caller. (**C**) Mutational signatures identified by non-negative matrix factorization (NMF) in the 15 Hupki MEF-derived clones (sig A, sig B, and sig C). X-axis represents the trinucleotide sequence context. Y-axis represents the frequency distribution of the mutations. The predominant trinucleotide context for T:A > A:T mutations is indicated in sig B (5’-C<u>T</u>G-3’). The trinucleotide contexts for C:G > G:C (5’-G<u>C</u>C-3’) and T:A > G:C mutations (5’-N<u>T</u>T-3’) are highlighted in sig C. (**D**) Contribution of the identified signatures to each sample (X-axis), assigned either by absolute SBS counts or by proportion (bar graphs). The reconstruction accuracy of the identified mutational signatures in individual samples is shown in the bottom scatter plot (Y-axis value of 1 = 100% accuracy).

### Variant allele frequency analysis

Variant allele frequency (VAF) analysis was carried out for GA clones. Overall, a significant proportion of acquired mutations was present at allelic frequencies between 25-75% (Suppl. Fig. S5). Upon grouping of substitutions into bins of high (67-100%), medium (34-66%) and low (0-33%) VAF, the predominant GA-specific mutation types (T:A>A:T, T:A>C:G and C:G>A:T) started manifesting at high VAF, whereas the 5’-NTT-3’ alterations, corresponding to the COSMIC signature 17 previously reported to arise in cultured mouse cells including MEFs (38,54,55) showed lower VAF, therefore a later appearance in the cultures (Suppl. Fig. S6). This observation suggests the early effects of the GA exposure and the reproducible contribution of the induced mutations to the senescence bypass and their clonal propagation during the immortalization stage.

### Mutational signature analysis

Using NMF, we extracted the mutational signatures from all the MEF clones. Using computed statistics for estimating the number of signatures, three signatures were identified as an optimal number, with signatures A and C enriched in the Spont and ACR clones, and signature B selectively enriched in the GA clones (Fig. 2C,D). Reconstruction of the observed mutation spectra supports the robustness of the signature analysis with strong Pearson’s correlation and cosine similarity in GA-derived clones (Fig. 2D). In signature C and also to a lesser extent in signatures A and B, we observed an admixture of a pattern identical to the orphan COSMIC signature 17 (T:A>G:C in a 5’-NTT-3’ trinucleotide context), described in various human cancers (most notably esophageal adenocarcinoma), but also seen in aflatoxin B1-driven mouse liver cancers (11), as well as primary MEF-derived clones (37,38). In *in vitro* contexts, this signature has been linked to cell culture conditions and associated oxidative stress (54,55). To refine further the obtained experimental signatures, we developed a signature ‘baiting’ approach that combined the MEF clones data with signature 17-rich data from esophageal adenocarcinomas from the ICGC ESAD-UK study for new NMF analysis (56). This resulted in considerable reduction (average = 47%, median = 48%) of the signature 17-specific most prominent T>G peaks and a more refined pattern for signature B, associated primarily with GA treatment (Fig. 3A and Suppl. Fig. S7). This putative GA signature retains the predominant enrichment for the T:A>A:T transversions and T:A>C:G transitions in the 5’-CTG-3’ and 5’-CTT-3’ trinucleotide contexts, and the C:G>A:T component. Moreover, these mutation types were marked by significant transcription strand bias (Fig. 3B and Suppl. Fig. S4), exhibiting higher accumulation of mutations on the non-transcribed strand consistent with the decreased efficiency of the transcription-coupled nucleotide excision repair due to adduct formation.

**Figure 3:**
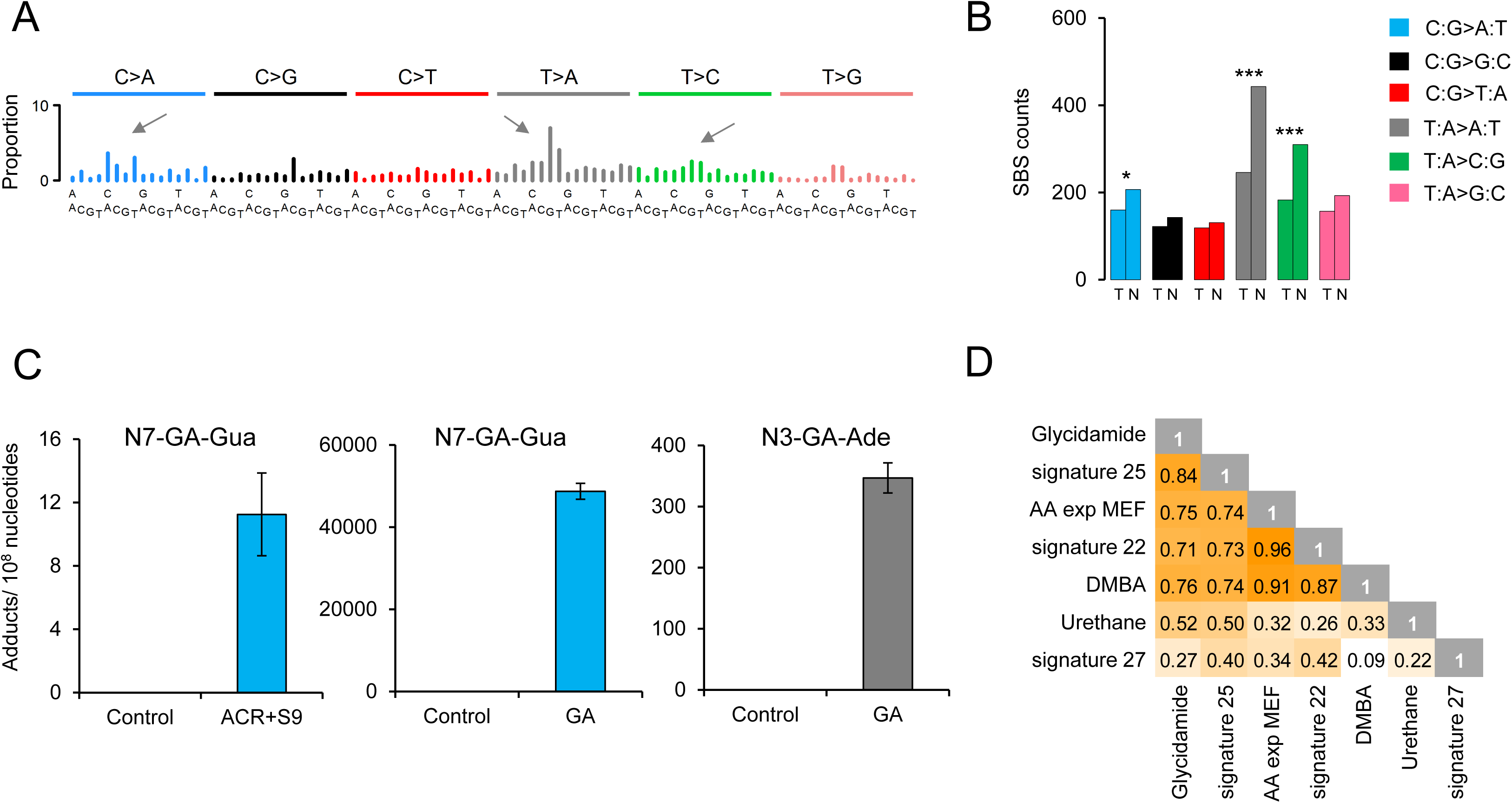
(**A**) Refinement of GA signature. The contribution of signature 17 (T:A>G:C in 5’-N<u>T</u>T-3’ context), present in all clones, was decreased by performing NMF on Hupki samples pooled with primary tumor samples with high levels of signature 17 (see Methods). (**B**) Transcription strand bias analysis for the six mutation types in GA-exposed clones. For each mutation type, the number of mutations occurring on the transcribed (T) and non-transcribed (N) strand is shown on the Y-axis. *** p < 10-^8^; * p < 10-^2^. (**C**) DNA adducts analysis as determined by LC-MS/MS. Levels of N7-GA-Gua adduct in ACR+S9 and GA treated MEFs and N3-GA-Ade DNA adduct level in GA treated MEFs. The data are presented as the number of adducts in 10^8^ nucleotides. *n **>** 2.* (**D**) Cosine similarity matrix comparing the putative glycidamide mutational signature with other A>T rich mutational signatures from COSMIC (signatures 22, 25, and 27) and from experimental exposure assays using specific carcinogens (7,12-dimethylbenz[a]anthracene (DMBA), urethane, and aristolochic acid (AA)).

### DNA adduct analysis

Following metabolic activation, acrylamide induces well-characterized glycidamide DNA adducts at the N7‐ and N3-positions of guanine and adenine, respectively. LC-MS/MS-based adduct quantification revealed the absence of these adducts in the spontaneously immortalized control samples as well as in MEFs exposed to acrylamide in the absence of S9 fraction (levels below the limit of detection). This suggests the lack of CYP2E1 activity, which is required for the metabolism of acrylamide to glycidamide, in the MEFs. Upon addition of human S9 fraction, N7-GA-Gua levels increased to 11adducts/10^8^ nucleotides, suggesting limited metabolic activation of acrylamide due to the presence of enzymatic activity in the S9 fraction (Fig. 3C and Suppl. Fig. S8). Glycidamide-exposed cells exhibited significantly increased DNA adduct levels, with both N7-GA-Gua and N3-GA-Ade observed at very high average levels, 49 000 adducts/10^8^ nucleotides and 350 adducts/10^8^ nucleotides, respectively, after subtracting the trace amount of contamination from the internal standard (Fig. 3C and Suppl. Fig. S8).

### Comparison of the glycidamide signature to known signatures characterized by prominent T:A>A:T profiles

We next performed cosine similarity analysis of the putative GA signature and all known T:A>A:T-rich signatures extracted from primary cancers as well as experimental systems (Fig. 3D and Suppl. Fig. S9). The best match was 84% pattern similarity with COSMIC signature 25 (derived from four Hodgkin lymphoma cell lines) (Fig. 3D). However, unlike the GA signature, COSMIC signature 25 exhibits strand bias for only T:A>A:T mutations and no transcription strand bias for the T:A>C:G mutations. Thus, the mutation patterns and strand bias on all three main mutation types generated by GA treatment (Fig. 3A,B) appear specific and novel.

### Glycidamide signature screening in human tumor data from the ICGC PCAWG

The initial mSigAct test performed on PCAWG data from lung and liver tumors indicated a marked presence of the GA signature. This observation was in keeping with the presence of acrylamide in tobacco smoke and was further corroborated by a cosine similarity of 94% between the adenine (T>N) components of COSMIC signature 4 (tobacco smoking) and the GA signature (Fig. 4A). We thus hypothesized that COSMIC signature 4 reflects co-exposure to B[a]P (generating C>N/guanine mutations with transcription strand bias) and to GA (generating T>N/adenine mutations with transcription strand bias) (Fig. 4A,B). To provide further experimental evidence, we generated a ‘pure’ B[a]P mutational signature by whole-genome sequencing of cell clones derived from B[a]P-exposed normal human mammary epithelial cells (HMEC). This yielded a robust signature characterized by predominant strand biased guanine (mainly C>A) mutation levels and negligibly mutated adenines (T>N) (Fig. 4A,B). Next, we used mSigAct to interrogate the PCAWG tumor samples for the level of exposure to the experimentally defined GA and B[a]P signatures (alongside other COSMIC mutational signatures) in 48 lung squamous carcinomas, 38 lung adenocarcinomas, and 320 liver cancers. We compared these to estimated levels of exposure to COSMIC signature 4, and found that in the lung cancers, a combination of the GA and B[a]P signatures accounted for very similar numbers of mutations as COSMIC signature 4, thus further supporting the hypothesis that COSMIC signature 4 represents combined and highly correlated exposure to GA and B[a]P (Fig. 4C). Compared to lung cancers, we found more variability in the assignment of mutation numbers to GA and B[a]P versus COSMIC signature 4 in liver cancers (Fig. 4C), which may reflect a decreased relationship between GA and B[a]P exposure due to generally more complex exposure history in the liver. The successful reconstruction of COSMIC signature 4 by the experimental GA‐ and B[a]p‐ signatures in the lung and liver human tumors enabled correct assignment of the GA-signature in a subset of 29 lung adenocarcinomas, 46 lung SCC and 26 liver tumors (Fig. 4D). The SBS counts corresponding to GA-mutational signature ranged between 300 up to 43,000 mutations/per sample in lung tumors, and between 190 to 23,000 mutations/per sample in liver tumors (Fig. 4D and Suppl. Table S5 – not available in the preprint version). These findings indicate exposure to glycidamide linked to tobacco smoking – when concomitant with B[a]P-signature, or through diet or occupation – in the absence of B[a]P signature (samples Liver-HCC::SP112224; Liver-HCC::SP49551; Liver-HCC::SP50105; Liver-HCC::SP98861; Liver-HCC::SP50183, see Suppl. Fig. S10 and Suppl. Table S5 – not available in the preprint version).

**Figure 4:**
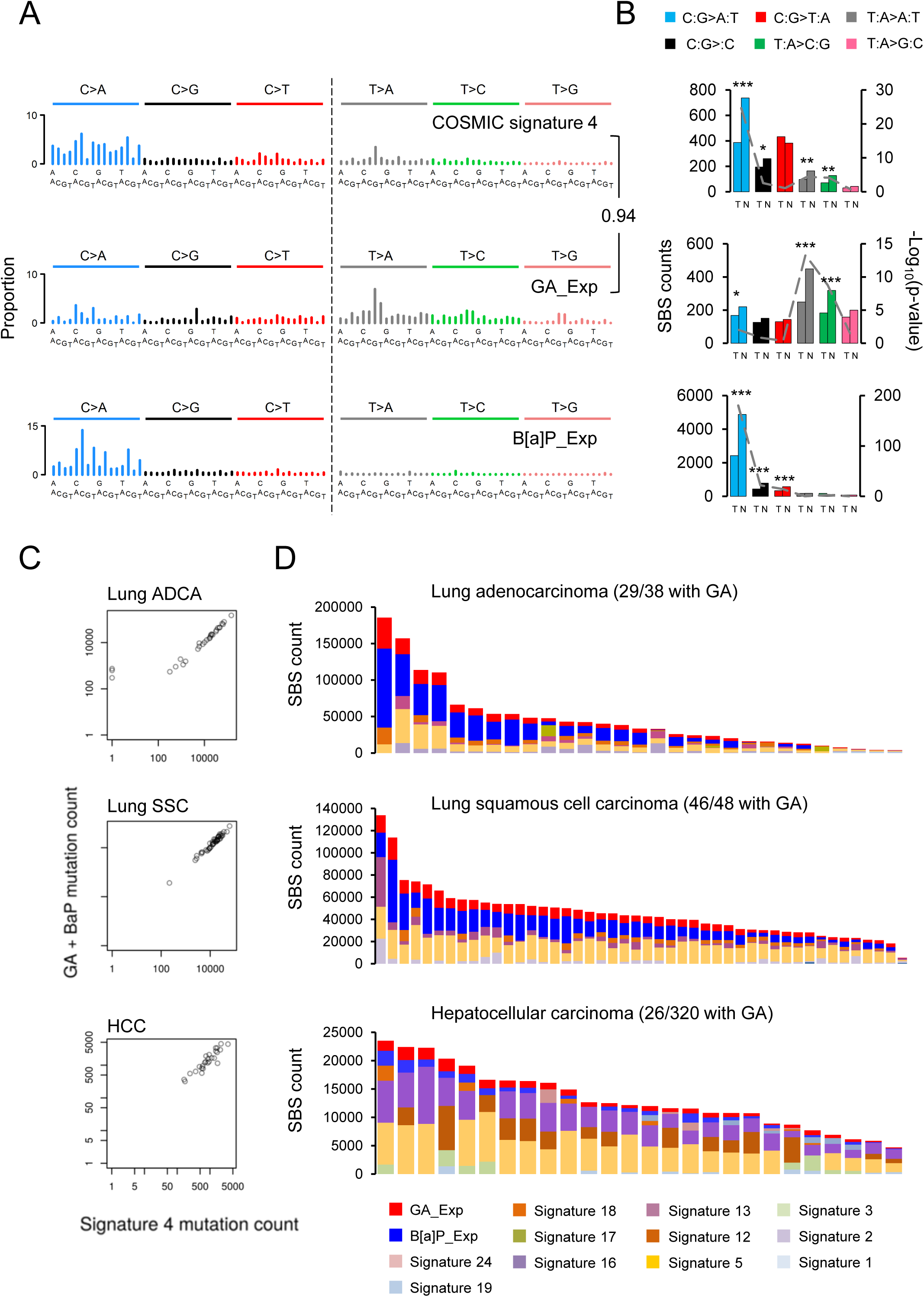
GA signature in human primary cancer genome PCAWG data. (**A**) Comparison of COSMIC signature 4 with two experimentally derived signatures (B[a]P_Exp = signature in clones from benzo[*a*]pyrene treated HMEC cells; GA_Exp = signature in clones from glycidamide-treated MEF cells). Cosine similarity between the T>N (adenine) components of signature 4 and GA signature is shown to the right. (**B**) Transcription strand bias analysis for the six mutation types underlying the signatures in panel A). For each mutation type, the number of mutations occurring on the transcribed (T) and non-transcribed (N) strand is shown on the left Y-axis. The significance is expressed as –log10(p-value) indicated on the right Y-axis. *** p < 10^-8^; ** p < 10^-4^; * p < 10^-2^. (**C**) Scatter plots show reconstruction of COSMIC signature 4 using B[a]p‐ and glycidamide‐ experimental mutational signatures in lung adenocarcinoma, lung squamous cell carcinoma and hepatocellular carcinoma from the PCAWG data set. (**D**) mSigAct analysis identifies the assignment and the contributions of mutational signatures (including the experimental signature_GA_Exp (red) and signature_B[a]P_Exp (blue)) to the mutation burden of a total of 101 PCAWG lung and liver tumors identified as positive for the GA signature signal.

## Discussion

In this study we report the identification of an exome-wide mutational signature for glycidamide, a metabolite of the probable human carcinogen acrylamide. The newly identified signature is based on massively parallel sequencing performed in a well-controlled experimental carcinogen exposure-clonal immortalization model, revealing characteristic mutagenic effects of glycidamide. The glycidamide mutational signature presented here and the results of statistical assessment of its presence in multiple human tumor types may help clarify the thus-far tenuous association of acrylamide with human cancer.

In concordance with its *in vivo* carcinogenicity in rodents (16,19,31,57), our findings in the established MEF carcinogen exposure and immortalization system suggest that characteristic mutagenic effects may play a role during acrylamide/glycidamide-driven tumor development. In contrast to glycidamide, acrylamide exposure led neither to an increased number of SBS nor did it induce characteristic mutation types in the MEF exposure system. Despite the absence of a mutagenic effect of acrylamide in our experiments, acrylamide and glycidamide exposures induce an almost identical set of tumors in both mice and rats, providing a substantial argument for a glycidamide-mediated tumorigenic effect of acrylamide (19). This is further supported by mechanistic studies showing that lung tissue from mice exposed to acrylamide and glycidamide displays comparable DNA adduct patterns as well as similar mutation frequencies in the *cII* transgene (36). Similar observations had been made in the context of *in vitro* mutagenicity of acrylamide in human and mouse cells, suggesting the key role for epoxide metabolite glycidamide to form pre-mutagenic DNA adducts (33).

As shown by our adduct analysis, acrylamide is not efficiently metabolized by MEFs. This finding is in keeping with the results from previous animal carcinogenicity studies. In fact, glycidamide induces hepatocellular carcinomas in neonatal B6C3F1 mice, whereas administration of acrylamide does not increase the tumor incidence. This has been attributed to the inability of neonatal mice to efficiently metabolize acrylamide (31). Moreover, in contrast to acrylamide treatment, glycidamide induces tumors of the small intestine in a dose-dependent manner upon perinatal exposure (57) and similar observations were made for glycidamide mutagenicity *in vitro* (33). We compensated for the lack of proper acrylamide metabolic activation by the addition of human S9 fraction, and the assessment of DNA adducts indeed suggests acrylamide metabolic activation upon addition of S9. However, the adduct levels are substantially lower compared to glycidamide exposure, which may account for the observed differences in mutagenicity. Interestingly, a consistent minor contribution of the glycidamide mutational signature was detected in the majority of ACR clones, whereas it was absent in the Spont clones. This raises the possibility that partial metabolic activation of acrylamide in the MEF system resulted in low levels of glycidamide. However, a clear mutational signature in the employed experimental setting was achieved only by exposing the cells directly to glycidamide.

Single reporter gene studies had previously linked acrylamide and glycidamide exposure to multiple different mutation types. Thanks to the larger number of mutations captured by exome sequencing, we were able to attribute to the glycidamide exposure a particular mutational signature characterized by strand-biased C:G>A:T and T:A>A:T transversions, and T:A>C:A transitions towards the non-transcribed strand suggesting a formation of DNA-adducts. The presence of N7-GA-Gua and N3-GA-Ade, two well-characterized glycidamide DNA adducts originating from the metabolic conversion of acrylamide (30,44,53), shows a remarkable relationship between DNA adduct profiles and the putative mutational signature of glycidamide. N3-GA-Ade and N7-GA-Gua are depurinating adducts. They can result in apurinic/apyrimidinic sites, which, during replication, induce the mis-incorporation of deoxyadenine, leading to the observed T:A>A:T and C:G>A:T transversions of the glycidamide signature, respectively. The third mutation type specifically enriched in the glycidamide signature, T:A>C:G transitions, has been ascribed to the N1-GA-Ade adduct, a miscoding adduct and the most commonly identified adenine adduct *in vitro* (35,44,53,58). Levels of the guanine adduct were especially high in the exposed MEF cells, whereas the associated C:G>A:T transversions in the resulting post-senescence clones were less represented. This could reflect differences in DNA repair efficiency concerning individual GA-DNA adduct species, or the fact that the resulting clones are derived from single cells whereas the GA-DNA adducts were measured on average in the bulk primary cell population. A mechanism of negative selection of cells with high N7-GA-Gua adduct burden is also plausible.

We observed consistent presence of COSMIC signature 17 in the data generated from the untreated and treated MEF clones. The etiology of signature 17 remains unknown. While some candidate causal factors have been proposed in esophageal adenocarcinoma and gastric cancers (e.g., inflammatory conditions due to acid reflux, *H. pylori*) (56) and in cultured mouse cell systems (54,55), further studies are required to establish why signature 17 tends to arise *in vitro* in immortalized clones derived from mouse embryonic fibroblasts as observed in our study and also previous work (38).

Genome-scale sequencing of tumor tissues will be needed to verify, *in vivo*, the glycidamide mutational signature identified in this study. The established animal models (18,19) of acrylamide‐ and glycidamide-mediated tumorigenesis provide a suitable starting point, and it would be interesting to compare mutational signatures derived from these models with the *in vitro* results. The identified glycidamide signature with its extended features of transcription strand bias for the major mutation types differs from the currently known COSMIC signatures (Fig. 3D). In addition, we show that in the cancer genome sequencing data sets from the ICGC PCAWG effort, the putative glycidamide-mutational signature can be identified in a subset of tumors of the lung and liver (sites of possible acrylamide exposure due to tobacco smoking), based on combining experimentally derived signatures with sophisticated computational signature reconstruction approaches (Fig. 4).

The continued interest in understanding the contribution of acrylamide and its electrophilic metabolite glycidamide to cancer development reflects recent accumulation of new mechanistic data on the animal carcinogenicity of the compounds. The possible carcinogenic effects in humans have been recommended for re-evaluation by the Advisory Group to the Monographs Program of the International Agency for Research on Cancer (59). Our findings related to the reconstruction of COSMIC signature 4 using the experimental GA-signature and B[a]P signature, together with the presence of the GA signature in the lung and liver cancer data are relevant given the established high contents of acrylamide in tobacco smoke. Despite the absence of prominent T>N (adenine) mutations in the experimental B[a]P exposure setting, we cannot exclude a possibility that in the human lung cells the adenine residues can be additionally targeted by other tobacco carcinogens such as benzo[*a*]pyrene derivatives or nitrosamines. Importantly, five liver tumor samples identified in this study harbored the GA signature but the major features of signature 4 as represented by the experimental B[a]P signature were absent (Suppl. Fig. S10, Suppl. Table S5 – not available in the preprint version). These tumors are thus of particular interest as they could reflect dietary or occupational exposure to acrylamide.

The presented mutational signature of glycidamide and its potential use for screening of cancer genome sequencing data may provide a basis for relevant assessment of cancer risk through new carefully designed molecular cancer epidemiology studies. Future validation analyses involving e.g. GA-DNA adduct monitoring in non-tumor tissue of cancer patients or in animal exposure models are warranted to provide additional evidence that the predominant T>N mutations in the cancers identified in this study indeed originate from exposure to acrylamide and its reactive metabolite glycidamide.

## Acknowledgments

The views expressed in this manuscript do not necessarily represent those of the U.S. Food and Drug Administration. The study was supported by funding obtained from INCa-INSERM (Plan Cancer 2015 grant to J.Z.), NIH/NIEHS (1R03ES025023-01A1 grant to M.O.), and the Singapore National Medical Research Council (NMRC/CIRG/1422/2015 grant to S.G.R.) and the Singapore Ministry of Health via the Duke-NUS Signature Research Programmes to S.G.R.. M.R.S. was supported by the U.S. Department of Energy under Contract No. DE-AC02-05CH11231. We thank the NYU Genome Technology Center, funded in part by the NIH/NCI Cancer Center Support Grant P30CA016087, and GENEWIZ, South Plainfield, NJ, USA, for expert assistance with Illumina sequencing.

